# Subtherapeutic doses of vancomycin synergize with bacteriophages for treatment of experimental methicillin-resistant *Staphylococcus aureus* infective endocarditis

**DOI:** 10.1101/2022.06.02.494631

**Authors:** Jonathan Save, Yok-Ai Que, José M. Entenza, Grégory Resch

## Abstract

**Background:** Recurrent therapeutic failures reported for treatment of methicillin-resistant *Staphylococcus aureus* (MRSA) infective endocarditis (IE) with vancomycin may be due to poor bactericidal activity. Alternative antibacterial approaches using bacteriophages may overcome this limitation.

**Objective:** An experimental rat model of MRSA IE (EE) was used to examine the efficacy of vancomycin combined with a 1:1 bacteriophage cocktail composed of *Herelleviridae* vB_SauH_2002 and *Podoviridae* 66.

**Methods:** Six hours after inoculation with ca. 5 log_10_ CFU of MRSA strain AW7, animals were treated with either: (i) saline, (ii) an equimolar two-phage cocktail (bolus of 1 mL followed by a 0.3 mL/h continuous infusion of 10 log_10_PFU/mL phage suspension), (iii) vancomycin (at a dose mimicking the kinetics in humans of 0.5 g b.i.d.), or (iv) a combination of both. Bacterial loads in vegetations, and phage loads in vegetations, liver, kidney, spleen, and blood, were measured outcomes.

**Results:** Phage cocktail alone was unable to control growth of strain AW7 in cardiac vegetations. However, when combined with subtherapeutic doses of vancomycin, a statistically significant decrease of Δ4.05 ± 0.94 log_10_ CFU/g at 24 h compared to placebo was detected (p < 0.001). Administration of vancomycin was found to significantly impact local concentrations of phages in the vegetations and in the organs examined.

**Conclusions:** Lytic bacteriophages as an adjunct treatment to standard of care antibiotics could potentially improve management of MRSA IE. Further studies are needed to investigate the impact of antibiotics on phage replication *in vivo*.

## Introduction

*Staphylococcus aureus* (*S. aureus*) endocarditis continues to be associated with substantial morbidity and mortality rates [1], especially when caused by methicillin-resistant strains (MRSA) [2]. Current guidelines recommend use of vancomycin for MRSA infective endocarditis (IE), while daptomycin is a reasonable alternative [3]. However, minimal penetration into vegetation [4], a slow bactericidal effect [5], and potentially severe side effects such as nephrotoxicity challenge this dosing scheme. Clinical failures associated with administration of vancomycin have included renal failure [6] and the emergence of vancomycin-intermediate *S. aureus* (VISA) clones [7]. Thus, alternative and/or adjunct treatments for MRSA IE are needed.

Therapeutic use of bacterial viruses called bacteriophages (phages), referred to as phage therapy, has been successfully applied to cases involving bacterial infections for nearly a century in several Eastern European countries [8]. Since the early 2000s, phage therapy has undergone greater investigation in Western societies, including for the treatment of *S. aureus* IE [9, 10]. Renewed interest for this forgotten cure is principally in response to a growing threat of antibiotic resistance [11]. Recently, however, a few randomized clinical trials have repeatedly failed to report phage therapy efficacy [12].

In sharp contrast to most experimental settings, phage have mainly been evaluated in conjunction with antibiotics [13]. However, use of antibiotics may represent a double-edged sword if it reduces the number of bacteria needed for phages to replicate. Recently, we reported a synergistic bactericidal effect of a novel anti-*S. aureus* two-phage cocktail in combination with the beta-lactam, flucloxacillin, for treatment of methicillin-susceptible *S. aureus* (MSSA) EE. However, the same study also observed that in the presence of flucloxacillin, the pharmacokinetic (PK) profile of the phages was significantly altered *in vivo* [14]. Here, we further investigate whether the same phage cocktail will synergize with the glycopeptide, vancomycin, for treatment of MRSA EE. We also evaluate to what extent vancomycin affects the PK profile of phages in this well-established rat model of EE [15, 16].

## Materials and methods

### Bacterial strains, bacterial genomic DNA, and bacteriophages

MRSA strain AW7 [17] was isolated from a patient in Switzerland with bacteraemia and has been used to establish EE [18] and pneumonia [19-22]. We adapted a protocol from Bae et al. [23] to obtain purified *S. aureus* genomic DNA (described in Supplemental Material). *Herelleviridae* phage vB_SauH_2002 was isolated from sewage water [14] and *Podoviridae* phage 66 was purchased from the National Collection of Types Cultures of Public Health England (catalogue #8289). These two phages were produced on MSSA Laus102 strain [24] and the concentration of each phage solution was adjusted to 10 log_10_plaque-forming units (PFU)/mL. Details regarding growth conditions and reagents are provided in Supplemental Material.

### *In vitro* activity of phages and vancomycin against planktonic cultures and biofilms

The lytic activity of each single phage and their combination as a 1:1 cocktail was evaluated in diluted drop tests, turbidity assays, or time-kill assays on planktonic AW7 cultures in the presence or absence of vancomycin at 1× and 10× minimum inhibitory concentration (MIC) [25]. A detailed description of the methods used is provided in the Supplemental Material section.

Synergy between both antibacterial treatments was defined as an additional ≥ 2 log_10_ colony forming unit (CFU)/mL reduction in bacterial load compared to the effect of the best single treatment [26].

### Murine model of EE

Female Wistar rats (Charles River, L’Abresle, France) weighing 180–200 g were used in this study. Animals were kept in specific pathogen-free rooms (12 h light / dark conditions, 23 ± 1 °C, water, and food *ad libitum*). The experiments involving animals were performed according to Swiss Animal Protection Law guidelines and were approved by the Cantonal Committee on Animal Experiments of the State of Vaud (approval N° VD 879.10). Animals were anesthetized with an intraperitoneal administration of ketamine (Ketalar^®^, 75 mg/kg) and xylazine (Xylasol^®^, 0.5 mg/kg). An intraperitonal administration of buprenorphin (Temgesic^®^, 0.15 mg/kg) just prior to surgery served as an analgesic.

#### Induction of infection

Catheter-induced sterile aortic vegetations were produced in rats as previously described [16]. Antibacterial drug administration was performed according to a dosing schedule that mimics the kinetics of human IV antibiotic therapy [27, 28]. For this purpose, an IV line was placed via the jugular vein into the superior vena cava and connected to a programmable infusion pump (Pump 44; Harvard Apparatus, Inc., South Natick, MA, USA) [27]. Bacterial inocula were prepared from dilutions of mid-exponential phase cultures of MRSA strain AW7 (optical density at 600 nm (OD_600nm_) = 0.6, ∼8 log_10_ CFU/mL). Twenty-four hours after surgery, each animal received via IV 500 µL inoculum [5.11 ± 0.48 log_10_ CFU, corresponding to 10 times the 90% infectious dose [27]]. Inoculum size was confirmed by counting isolated colonies on trypticase soy agar plates (TSA; BD Difco™, Becton Dickinson, Sparks, MD, USA). Three uninfected animals were used for PK studies. Additional information related to the EE model used is provided in Supplemental Materials.

#### Power calculation

We hypothesized that 100% and 30% of the placebo and phage cocktail/vancomycin-treated rats would exhibit infected vegetations at 24 h, respectively. These estimates, along with an α = 0.05 and power (1-β) = 0.8, indicated that a sample size of at least eight animals per group was needed [29].

### Treatment Protocol

Six hours after bacterial challenge, a subset of animals was euthanized to assess the concentrations of bacteria present in vegetations at the onset of treatment. The remaining animals were subsequently treated with either: (i) mock therapy (saline), (i) a phage cocktail (10 log_10_ PFU/mL, equimolar concentration of each phage) injected as a 1-mL bolus followed by continuous infusion at 0.3 mL/h for up to 48 h, (ii) a subtherapeutic IV dose of vancomycin mimicking human kinetic treatment of 0.5 g given every 12 h for up to 48 h [30], or (iv) a combination of both treatments (Supplemental Figure 1). Programmable infusion pumps were used to deliver the treatments. Concentrations of bacteria in vegetations were assessed again 24 h and 48 h after the onset of treatment in the remaining rats that were euthanized. Concentrations were determined from colony counts of TSA plates. Details regarding the exact numbers of animals used in each group are reported in the Supplemental Material section.

### Outcomes

Primary outcomes included bacterial loads within cardiac vegetations at 24 h and 48 h after onset of treatment. Secondary outcomes included phage concentrations at 24 h and 48 h in cardiac vegetations, blood, spleen, liver, and kidneys. Selection of phage-resistant clones from the cardiac vegetations obtained under phage/vancomycin combinations were examined at 48 h. Methods used to assess all outcomes are described in Supplemental Material.

### Statistical analysis

Data obtained from time-kill assays and from the *in vivo* model of IE were compared by using two-way analysis of variance (ANOVA) with Tukey’s multiple comparisons test. The Mann-Whitney test was used to compare PK data from healthy and MRSA-infected animals. All statistical analyses were performed with GraphPad Prism software (version 9.0.0, La Jolla, CA, USA). Statistical test results were considered significant when p ≤ 0.05 were obtained. Mean values are reported with standard deviation (SD).

## Results

### *In vitro* activity of vB_SauH_2002/phage 66 cocktail on MRSA AW7

A phage cocktail (with a non-diluted phage titer of 10 log_10_ PFU/mL) composed of an equimolar mixture of *Herelleviridae* vB_SauH_2002 and *Podoviridae* 66 formed clear plaques on a lawn of MRSA strain AW7 in a diluted drop test assay (Figure 1A). In turbidity assays, a MOI of at least 0.1 was needed to prevent growth of strain AW7 over 24 h (Figure 1B). In time-kill assays, the phage cocktail at an MOI of 1 was highly bactericidal at 4 h (Δ3.09 ± 0.80 log_10_ CFU/mL vs. onset of treatment; p < 0.01). However, regrowth was observed at 24 h (Figure 1C), and was prevented with addition of 2x MIC vancomycin. Phage/vancomycin treatment also exhibited synergistic activity compared to vancomycin alone and achieved an additional Δ2.21 ± 0.75 log_10_ CFU/mL reduction at 24 h (Figure 1C, p < 0.05).

**Figure 1.**
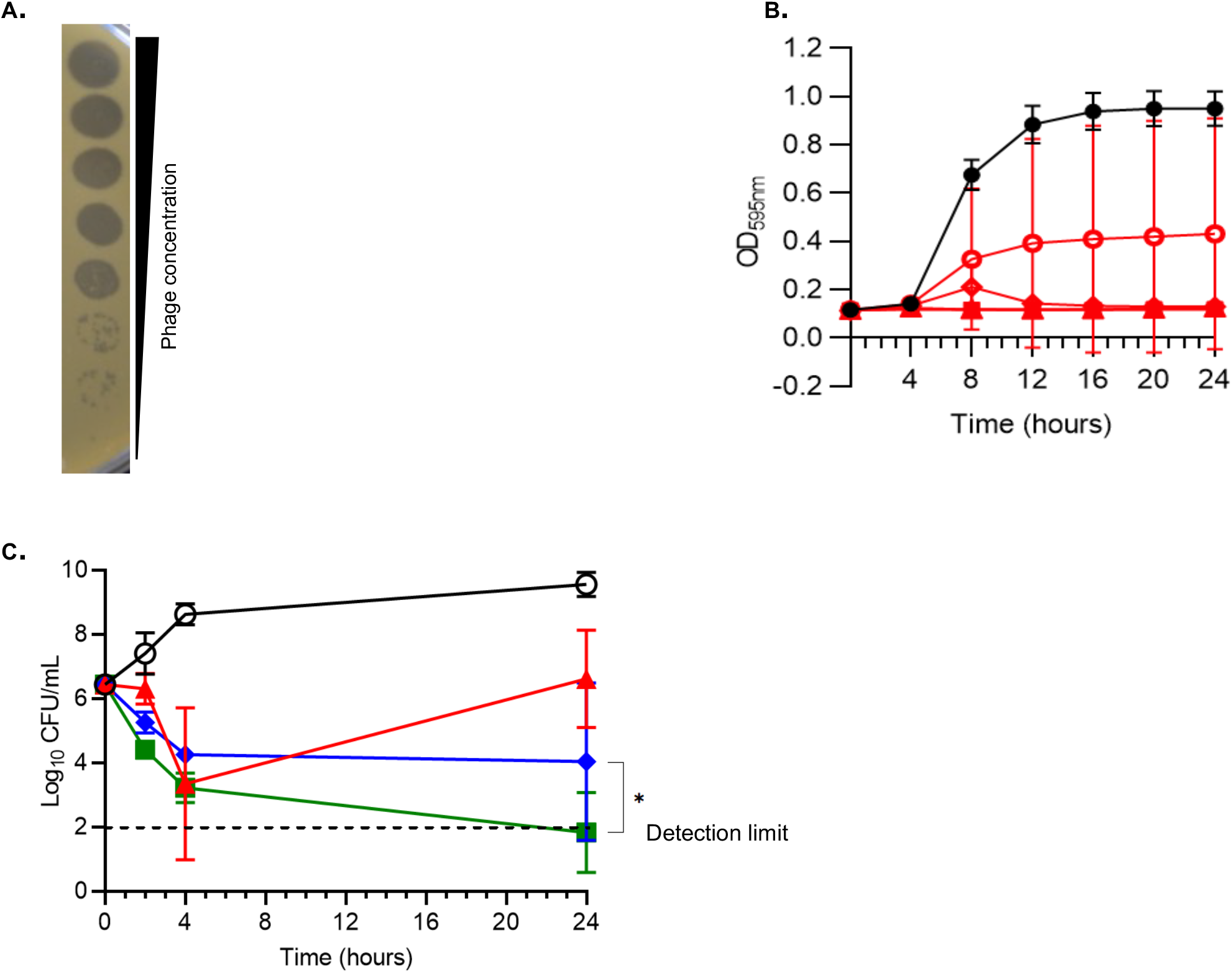
*In vitro* activity of the phage cocktail against MRSA AW7. **A.** Diluted drop tests, **B.** Turbidity assays, **C.** Time-kill assays in absence and presence of vancomycin. **A.** The phage cocktail was serially 10-time diluted from top to bottom (starting concentration was 10^10^ PFU/mL). **B.** Control without phages (closed black circles); phages at MOI = 0.01 (open red circles), MOI = 0.1 (open red diamonds), MOI = 1 (open red triangles), MOI = 10 (closed red squares), and MOI=100 (closed red triangles). **G.** Time-kill assays were performed by challenging MRSA AW7 with saline (open black circles); phage cocktail at MOI = 1 (closed red triangles), vancomycin at 2xMIC (closed blue diamonds) or a combination of both treatments (closed green squares). Means (±SDs) of three independent experiments performed in triplicate are shown in panels B and C. One-way ANOVA with Tukey correction for multiple comparison statistical tests were performed to compare either areas under the curves of curves obtained at MOI = 0.1 (Figures 1B) or 24 h’ time-points (Figure 1C). CFU, colony forming unit; MOI, multiplicity of infection; PFU, plaque forming unit.

### Subtherapeutic doses of vancomycin synergize with the phage cocktail *in vivo*

Without treatment, bacterial loads within cardiac vegetations rose from 6.55 ± 0.59 log_10_ CFU/g at the onset of therapy (i.e., 6 h post-inoculation) to 9.99 ± 0.70 log_10_ CFU/g after 24 h (Figure 2B, p < 0.01). Administration of either the phage cocktail or subtherapeutic doses of vancomycin failed to control bacterial growth within the vegetations at 24 h (10.55 ± 0.60 log_10_ CFU/g vs. 8.53 ± 1.65 log_10_ CFU/g, respectively) compared to placebo (9.99 ± 0.70 log_10_ CFU/g) (p = 0.99 and p = 0.59, respectively). When the phage cocktail was administered in combination with subtherapeutic doses of vancomycin, a significant decrease in bacterial load was observed within cardiac vegetations at 24 h compared to placebo (Δ4.05 ± 0.94 log_10_ CFU/g, p < 0.001). This decrease in bacterial load was also superior to that achieved with a subtherapeutic dose of vancomycin alone (5.93 ± 2.56 log_10_ CFU/g vs. 8.53 ± 1.65 log_10_ CFU/g, respectively; p < 0.05). Furthermore, phage and vancomycin exhibited synergistic activity to produce a Δ2.60 ± 1.0 log_10_ CFU/g decrease in bacteria load compared to antibiotic alone. The latter combination remained bacteriostatic at 24 h compared to the onset of therapy (5.93 ± 2.56 log_10_ CFU/g vs. 6.55 ± 0.59 log_10_ CFU/g, respectively; p = 0.27). Maximal bacterial killing was achieved when the phage/vancomycin combination was administered for 48 h (Δ1.98 ± 0.67 log_10_ CFU/g decrease compared to the onset of treatment; p = 0.051) (Figure 2B).

**Figure 2.**
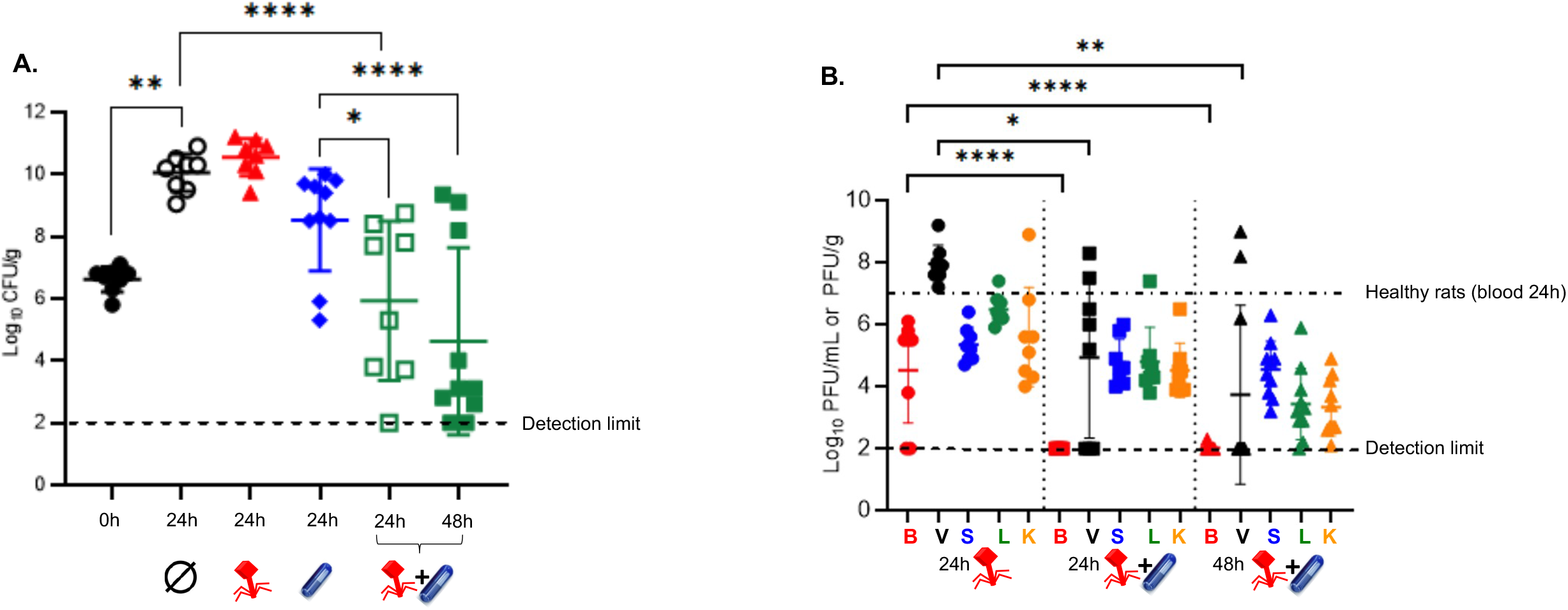
Treatment of EE with the phage cocktail in the presence or absence of a subtherapeutic dose of vancomycin. **A.** Bacterial loads in cardiac vegetations measured at 6 h post infection (i.e 0 h or onset of treatment) in the control rats (closed black circles, n = 8) and 24 h after the onset of treatment in rats given a mock therapy (saline, open black circles, n = 8), the phage cocktail alone for 24 h (closed red triangles, n = 8), a low dose of vancomycin alone for 24 h (closed blue diamonds, n = 10), or the phage cocktail in combination with vancomycin for 24 h (open green squares, n = 8) and 48 h (closed green squares, n = 10). Each symbol represents one animal. **B.** *In vivo* phage pharmacokinetics. Phage concentrations in blood (red), cardiac vegetations (black), spleen (blue), liver (green), and kidneys (orange) from rats 24 h after initiation of treatments with either the phage cocktail alone (left; n = 8) or the phage cocktail and vancomycin combination (middle; n = 8) and 48 h after initiation of treatment with the phage cocktail and vancomycin combination (right; n = 10). The black dotted-dashed line represents the mean concentration of phages at 24 hours in the blood of healthy rats treated with the phage cocktail (alone) (n = 3). Means (±SDs) are reported. \**p* < 0.05; \*\**p* < 0.01; \*\*\*\**p* < 0.0001; one-way ANOVA with Tukey’s multiple comparisons test.

### Subtherapeutic doses of vancomycin impair the PK profile of phage *in vivo*

The mean phage concentration determined from the blood of infected rats after a 24-h IV infusion of the phage cocktail did not statistically differ from that determined from the blood of healthy rats (4.53 ± 1.70 log_10_ PFU/mL vs. 6.90 ± 0.41 log_10_ PFU/mL, respectively; p = 0.47) (Figure 2C). In contrast, subtherapeutic doses of vancomycin strongly affected the PK profiles of phages in the blood and across the organs tested. For example, there were very few phages recovered from the blood in 1 out of 10 rats 48 h after phage-vancomycin combined therapy (2.02 ± 0.08 log_10_ PFU/mL vs. 4.53 ± 1.70 log_10_ PFU/mL for phages alone in infected animals, respectively; p < 0.0001).

Furthermore, glycopeptide treatment decreased phage availability within cardiac vegetations by Δ3.03 ± 1.16 log_10_ PFU/g at 24 h and by Δ4.33 ± 1.10log_10_ PFU/g at 48 h (p < 0.05 and p < 0.01, respectively) (Figure 2C).

### The phage 66-resistance phenotype observed *in vivo* is not linked to genetic determinants

The best treatment regimen (i.e. phage/vancomycin for 48h) in this study was not able to completely eradicate bacteria from cardiac vegetations. Therefore, we next investigated the possible role of resistance selection in the persistence of infection. None of the 53 single clones recovered from vegetations of the experimental rats treated with the combination therapy for 48 h displayed increased MIC for vancomycin (data not shown). There were six clones out of 53, which developed resistance to phage 66, yet not to phage vB_SauH_2002 or the phage cocktail (SRS pattern, Supplemental Table 1 and Supplemental Figure 2B). Using clone 17C4-SSS as a reference genome, comparative genomics identified several non-synonymous mutations in five representative SSS clones and the six SRS clones. These mutations are located within coding genes or in intergenic regions (Supplemental Tables 2 and 3). Interestingly, the mutations observed in both the SSS and SRS clones exhibit marked similarity and are mainly located in genes coding for transposases of the IS4 family (Supplemental Tables 2 and 3). A single unique mutation (leading to Gly18Cys) was detected only in SRS clone C11 in gene 993 that codes for an IS4 transposase (Supplemental Tables 2 and 3, highlighted in red). However, this same transposase was also found impacted by two other non-synonymous mutations (leading to Val41Gly and Gly20Asp) present in both the SSS and other SRS clones (Supplemental Tables 2 and 3).

### Despite efficient infection, phage 66 is not able to control *in vitro* growth of planktonic MRSA AW7 cells

Since phage 66 was found more prone to resistance development than phage vB_SauH_2002, we further characterized its *in vitro* bactericidal activity against MRSA AW7. In double layer assays, phage 66 formed PFU on a lawn of MRSA AW7. Moreover, the PFU were more turbid than the PFU observed for phage vB_SauH_2002 alone, or for the phage cocktail (Supplemental Figure 2A). Interestingly, the PFUs were devoid of surrounding halos, which were previously reported on MSSA Laus102 [14]. Finally, compared to phage vB_SauH_2002, which was able to control growth of planktonic MRSA AW7 cells *in vitro* for 24 h at an MOI = 0.1 (Supplemental Figure 3A), phage 66 was ineffective even at MOI = 100 (Supplemental Figure 3B).

## Discussion

We recently developed a promising two-phage cocktail that strongly synergizes with the standard of care antibiotic, flucloxacillin, in the treatment of MSSA EE [14]. Although the two-phage cocktail displayed very close *in vitro* activity in diluted drop tests and turbidity assays against MRSA strain AW7 as MSSA strain Laus102, it could not control bacterial growth in our *in vivo* rat model of MRSA EE. It has been demonstrated that predicting *in vivo* results based on *in vitro* testing is a difficult task in the context of phage therapy [31]. In the present study, a decrease of Δ3.09 ± 0.80 log_10_ CFU/mL in 4 h was observed for MRSA AW7. Previously, a decrease of Δ4 log_10_ CFU/mL in 2 h was observed for MSSA Laus102 [14]. Accordingly, a delayed bactericidal effect in time-kill assays was observed for MRSA AW7 in the present study, and this result is consistent with the total lack of efficacy observed for the phage cocktail alone against AW7 in our *in vivo* model.

In addition, therapeutic differences observed for the phage/antibiotic combination between MRSA and MSSA isolates in animals with EE may partially be due to impairment of phage PK profiles by vancomycin. It is generally acknowledged that a virulent phage infection leads to a high phage burst (referred to as auto-dosing), and this is a pre-requisite for the efficacy of phage therapy *in vivo* [32]. Our results further support this assumption, as the concentration of phages within cardiac vegetations was 2 log_10_ PFU/g lower in the rats infected with MRSA AW7 compared to the animals infected with MSSA Laus102. Even greater reductions in phage titers (i.e., ranging from Δ3 to Δ4.5 log_10_PFU/g) were observed across the organs that were evaluated compared to the levels previously measured with MSSA Laus102 [14]. As a result, MOI of 0.001 achieved within the cardiac vegetations in the MRSA AW7 setting was much lower than the MOI of 100 observed with MSSA Laus102 [14]. This result potentially contributed to the observed lack of efficacy of phages alone against MRSA AW7. Indeed, in the turbidity assays, lack of AW7 growth control by the phage cocktail was already observed at MOI 0.01. Whether improvement of the phage PK profile would have resulted in higher *in situ* MOIs, and subsequently a better bactericidal effect against the MRSA strain, remains to be evaluated. However, the latter could prove challenging to investigate due to current limitations associated with the production of highly concentrated batches of both phages.

The results of the present study confirm that a combined phage/antibiotic treatment regimen has the potential to outperform single treatment [12]. Yet, a lower efficacy of the phage component against MRSA AW7 eventually translated into a significantly lower therapeutic efficacy of the phage/antibiotic combination against MRSA AW7 compared to MSSA Laus102. Bactericidal activity was not achieved with the phage/vancomycin combination treatment, even after 48 h. In contrast, the phage/flucloxacillin combination previously exhibited strong bactericidal activity with a Δ5.25 log10 CFU/g decrease in bacteria in vegetations after 24 h [14].

Addition of antibiotic strongly affected the PK profile of the phages tested. A negative impact on their auto-dosing capacity was observed [33]. Moreover, even a subtherapeutic dose of vancomycin dramatically reduced the already very low capacity of the phage to replicate *in vivo* on MRSA AW7. As a result, the concentration of phage reached at 24 h in the vegetations infected with MRSA AW7 was ca. Δ4 log10 PFU/g lower than the concentration reached following infection with MSSA Laus102 under flucloxacillin therapy [14]. There were also no viable phage particles recovered at 24 h from the blood of the MRSA-infected animals, while 4.52 ± 1.70 PFU/mL was previously detected in the blood of animals treated with a combination of phage and flucloxacillin [14]. It remains to be determined whether this result is due to phage/bacteria interactions and/or a class effect of the antibiotic present. In addition, variables regarding the modes of administration of phage and antibiotic should be evaluated. For example, sequential administration of treatments, which has only occasionally been studied to date [34], could be investigated.

Based on the present results, careful adjustment/personalization of phage cocktail compositions according to the infecting strain, and their use with or without antibiotics, warrants further study. Indeed, our investigation of the activity of both types of phages revealed major differences in their antibacterial activity, particularly for phage 66. On MRSA AW7, turbid lysis was observed in diluted drop tests, and an absence of depolymerase halos was noted around phage 66 PFUs compared to MSSA Laus102 [14]. Moreover, while phage 66 efficiently prevented *in vitro* growth of Laus102 for 16 h at low MOIs [14], it was not able to control AW7 *in vitro* growth even at MOI of 100. These results may have contributed to the unfavourable PK profile for the phage cocktail tested in rats infected with MRSA AW7. In future studies, the concentration of each phage needs to be measured separately.

It has been reported that a subset of Laus102 phage 66-resistant variants carry mutations in the *tarS* coding region of a glycosyltransferase which mediates beta-O-N-acetylglucosaminylation of wall-teichoic acid (WTA) [14]. However, we did not identify any AW7 phage 66-resistant variants carrying mutations in this region. This was surprising since *tarS* mutations, which can result in removal of beta-O-N-acetylglucosaminylated WTA from the cell wall, have been characterized as a major mechanism of resistance to *Podoviridae* [35, 36]. Furthermore, no specific genetic determinants were found to be associated with *in vivo* development of resistance to phage 66 in MRSA AW7. While it is possible that such clones could have been lost during sample processing and/or storage, our observations suggest that resistance to phage 66 is driven by an adaptive phenotypic plasticity in AW7 rather than by genetic mutations. Indeed, it has been demonstrated that variations in bacteriophage activity can occur between different environments, and these are likely to be secondary to different glycosylation patterns of WTA [37, 38].

Thus, mechanisms of bacterial resistance to phage may have multiple pathways and components, even within the same bacterial species. Systematic studies of these mechanisms could provide greater insight into possible targets with the goal of preventing therapeutic failures.

## Acknowledgements

We thank Aurélie Marchet, Sabrina Pereira-Pipa, and Nathalie Bonvin for providing outstanding technical assistance.

